# Multidimensional dynamics of object representations in the human visual system

**DOI:** 10.64898/2026.04.27.720701

**Authors:** Zirui Chen, Leyla Isik, Michael F. Bonner

## Abstract

Natural image representations are distributed across many dimensions of visual cortex activity, but little is known about how the multidimensional structure of these representations evolves over time following stimulus onset. Here we examined the temporal dynamics and latent dimensional structure of natural object representations in large-scale EEG and MEG data. We also compared these data with leading representational models derived from large-scale human similarity judgments and deep neural networks. Our findings reveal a rapid expansion of stimulus dimensionality in the brain, which peaks within 100 milliseconds and gradually decays over hundreds of milliseconds. The dynamics of these dimensionality changes tracked the decoding accuracy for both behavioral embeddings and neural network features, suggesting that dimensionality may be a general indicator of representational expressivity. Interestingly, the dimensionality of the neural representations could not be fully explained by leading behavior-based or neural network models. Follow-up experiments showed that the remaining neural variance carried additional perceptually relevant information not yet explained by leading models. Together, these findings reveal previously unrecognized complexity in measurements of dynamic human brain responses to natural objects.

## 1 Introduction

The dimensionality of visual representations in the brain remains a fundamental question in neuroscience. One perspective suggests that the visual system implements dimensionality reduction on sensory inputs to achieve robust, low-dimensional representations of behaviorally relevant information [14, 33, 32, 39, 3]. This view has motivated researchers to study visual representations by focusing on low-dimensional embeddings of neural activity [11, 28, 31, 42]. However, advances in large-scale neural data collection [1, 25, 5] and analysis methods have shown that visual cortex representations of natural stimuli are not restricted to a low-dimensional subspace—instead they are scale-free, with stimulus-related information encoded along all available dimensions [15, 40, 22]. These findings suggest that conventional approaches focusing on leading dimensions may capture only a small fraction of the meaningful information encoded in cortical activity.

Despite these advances, the dynamics of multidimensional structure in human visual representations remain largely unexplored. Many previous studies of dynamics in human vision have examined feature decoding in electroencephalography (EEG) and magnetoencephalography (MEG) data and identified representational features with distinct temporal profiles, with more complex features tending to emerge at later time points [30, 4, 8, 9, 10, 20, 41, 19, 18, 36, 21]. However, what is not known is how the global structure of the representations evolves over time and how these global changes may be related to the dynamics of feature decoding.

A challenge of analyzing the multidimensional structure of EEG and MEG data is that these recording techniques suffer from signal spread across channels and lower signal-to-noise ratio compared with fMRI [37, 20]. Because of this, many EEG/MEG studies have employed dimensionality reduction as a preprocessing step [2, 4, 23]. However, in large-scale neural datasets, even dimensions with a low signal-to-noise ratio can carry reliable stimulus-related information and may be behaviorally relevant [15, 22]. Thus, by restricting analyses to leading dimensions and pre-determined features, previous studies may have overlooked potentially rich structure in the remaining neural dimensions.

Here we characterized the multidimensional structure of stimulus-related signal in human EEG and MEG responses to natural images. We examined two large-scale datasets containing responses to thousands of object images, and we compared neural representations with model features derived from behavioral similarity judgments and deep neural networks (DNNs) [16, 25, 24]. Our analyses show that the representational dimensionality of human brain responses rapidly increases after stimulus onset and remains high for hundreds of milliseconds. Conventional decoding analyses show that features derived from behavioral embeddings and DNNs are represented most strongly at times of peak neural dimensionality. However, further analyses using cross-decomposition show that these leading representational models do not fully explain the dimensionality of the neural signal. A follow-up behavioral experiment demonstrates that the remaining unexplained variance in neural responses contains perceptually relevant information. Together, these findings show that large-scale EEG and MEG data contain rich multidimensional structure that evolves rapidly after stimulus onset, carries behaviorally relevant information, and extends beyond what current representational theories capture.

## 2 Results

### 2.1 Time-resolved neural responses exhibit rich multidimensional structure

We examined the dynamics of representational dimensionality in human brain responses to natural images. To do so, we leveraged two large-scale datasets of time-resolved responses to thousands of images, THINGS-EEG2 and THINGS-MEG [16, 25] (Fig. 1a). The scale and richness of these datasets make them ideally suited for characterizing the multidimensional structure of visually evoked EEG and MEG responses. Both datasets contain responses to images of objects in their natural contexts, and they were sampled from the THINGS database [27].

**Figure 1:**
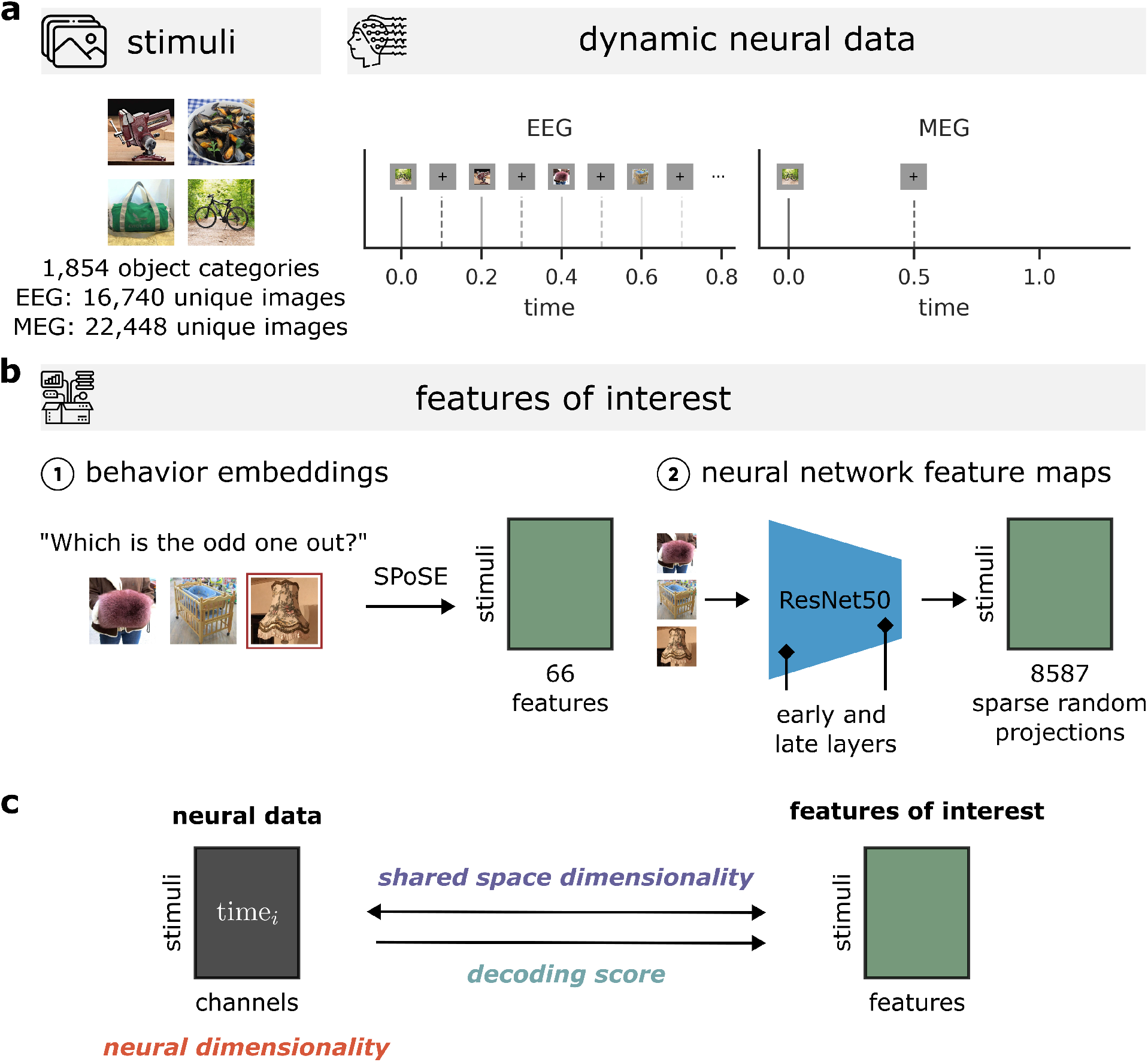
Overview of datasets and metrics evaluating neural representations and visual features of interest. **(a)** Left: Both EEG [16] and MEG [25] datasets are analyzed in this work and visual features of interest are vector representations related to these natural object images, with examples shown here. The stimuli are from the THINGS database, which contains images from 1,854 object categories [27]. Right: The two neural datasets have different experimental designs: the EEG dataset presented images for 100 millisecond (ms) and had 100 ms of fixation in between, while the MEG dataset presented images for 500 ms and had 1 s of fixation in between. **(b)** Left: Sparse nonnegative embeddings (SPoSE) were learned to explain human perceived similarity judgments in an odd-one-out behavioral experiment. Right: Features were extracted from layers of a ResNet50 model, trained on the YFCC15M dataset with the CLIP objective [38, 29]. Two layers were selected to capture low- and high-level visual features, and these DNN representations were reduced using sparse random projections. **(c)** We assessed the dimensionality of image representations at each individual time point. Further, to compare neural data and model features, we computed both shared-space dimensionality—a symmetric method using cross-decomposition [15, 22]—and decoding scores—a traditional asymmetric method using ridge regression.

THINGS-EEG2 contains EEG recordings in ten subjects who viewed 16,740 unique images, and THINGS-MEG contains MEG recordings in four subjects who viewed 22,448 unique images. The stimulus sets for both studies were designed with a training/test split. In the EEG dataset, the training set consists of ten unique images per 1,654 object categories, each shown four times, and the test set consists of one unique image from a different set of 200 object categories, each shown 80 times. In the MEG dataset, the training set consists of 12 unique images per 1,854 object categories, each shown once, and the test set consists of one unique image from a different set of 200 object categories, each shown 12 times. In both studies, subjects viewed the images while maintaining central fixation and performing an orthogonal target-detection task to ensure they were paying attention. Besides differing in imaging methods, these datasets also notably differ in their stimulus presentation timing, with the EEG dataset having much a shorter stimulus presentation time (100 ms for EEG and 500 ms for MEG) and a much shorter inter-stimulus interval (100 ms for EEG and 800-1,200 ms for MEG). The stark differences in the imaging methods and experimental paradigms of these datasets allow us to reveal dimensionality patterns that are either dependent on or invariant to experiment settings.

We characterized the dimensionality of stimulus representations in these data by applying principal component analysis (PCA) and estimating the reliability of the principal components (PCs) in held-out test data [15, 40, 22]. For each time point, we learned the PCs of multi-channel response patterns to all training images. We then projected the test-set responses onto these PC bases and evaluated the reliability of each PC by computing its correlation across split halves of the stimulus repetitions. We computed a summary statistic of representational dimensionality by counting the number of PCs with significant reliability (*p* < 0.05, cluster-based permutation test). We performed these analyses for all channels in the occipital and parietal cortices, which encompassed 17 channels in the EEG data and 80 channels in the MEG data. Computing this summary statistic at each time point allowed us to evaluate how the dimensionality of stimulus representations evolves over time.

The results are illustrated in Figure 2. For both datasets, they reveal a rapid increase in dimensionality after stimulus onset, peaking within 100 ms. Peak dimensionality is then sustained for hundreds of milliseconds (until 350 ms in the EEG data and 500 ms in the MEG data), followed by a gradual decay extending to nearly 800 ms. The similarities across the two datasets are notable given the major differences in their experimental paradigms. In the MEG experiment, the stimuli were displayed for 500 ms with a long interstimulus interval, allowing for the representations to evolve after stimulus onset without interference from other image inputs. However, even when stimuli are presented for a brief window of 100 ms and followed by a rapid succession of other sensory inputs, as in the EEG experiment, representational dimensionality nonetheless maintains peak levels for hundreds of milliseconds before gradually decaying. This suggests that the dimensionality dynamics observed here reflect a general property of image processing in the brain and are not strongly contingent on the details of stimulus presentation timing.

**Figure 2:**
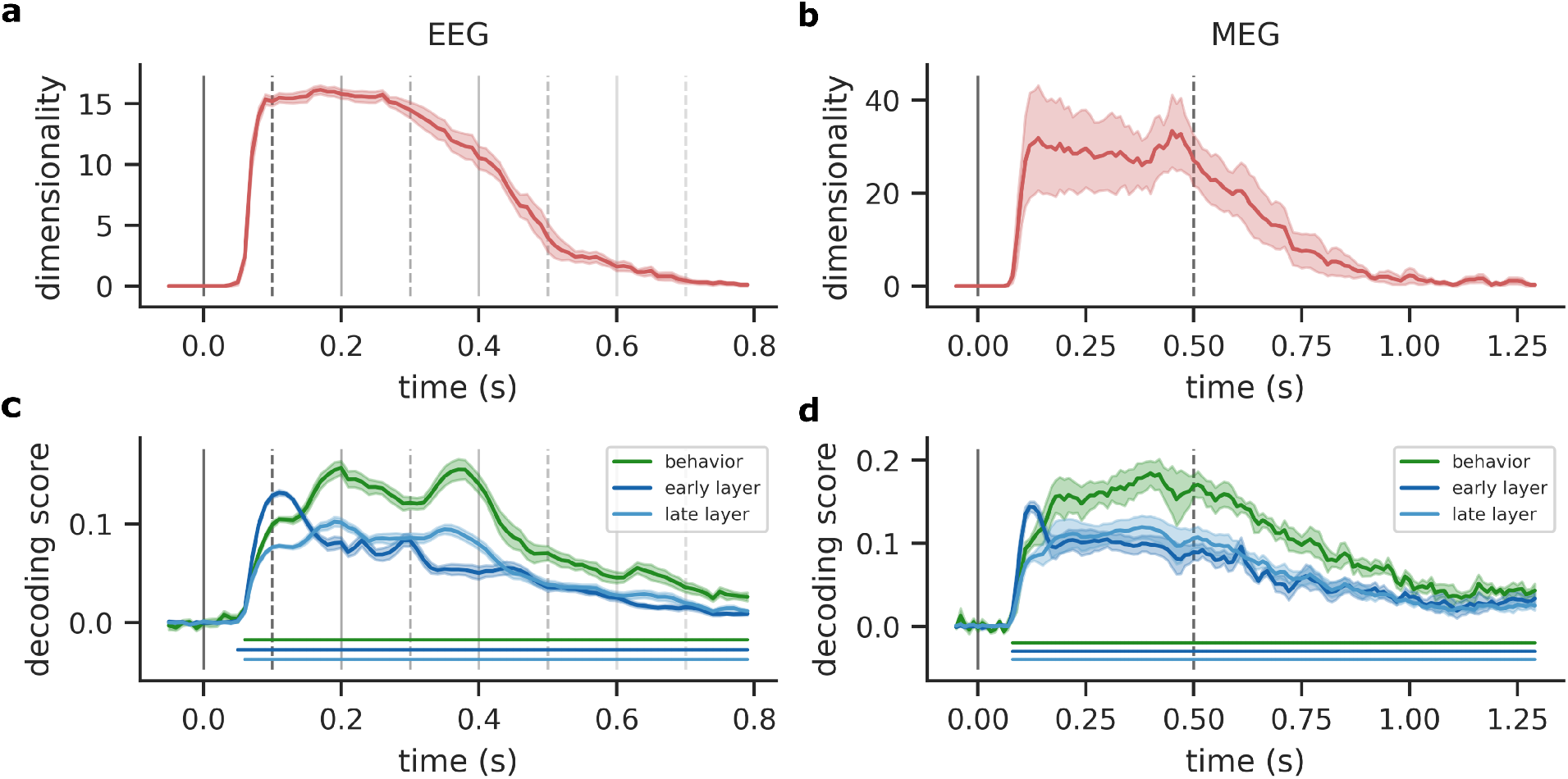
Temporal dynamics of neural dimensionality and feature decoding. **(a, b)** These plots show the dimensionality of stimulus-related variance in the EEG and MEG datasets at each time point following stimulus onset **(c,d)** These plots show the average decoding scores for behavioral embeddings and neural network features at each time point. The vertical lines show the stimulus-presentation timing of the two datasets, with solid lines indicating stimulus onsets and dashed lines indicating offsets. The error bars for dimensionality represent the standard error across subjects and those for decoding scores represent the standard error of feature-averaged means across subjects. For both datasets, the dimensionality increases sharply after stimulus onset and is sustained from 100-300 ms in the EEG data and 100-450 ms in the MEG data before gradually declining. All three sets of features can be decoded from the neural responses, and the decoding scores are generally highest during time points of high neural dimensionality. Horizontal lines under the decoding time courses indicate time windows of significant decoding (*p* < 0.05, cluster-based permutation test).

In a control analysis, we performed the same procedure in a region where we expected to find less stimulus-related signal and thus lower dimensionality (Fig. S1). Specifically, we examined the 21 channels from the frontal lobes in the EEG data and the 64 channels from the frontal lobes in the MEG data. As expected, the dimensionality of stimulus representations in the frontal lobes was low and transient. This contrasts with the findings in the occipital and parietal lobes, where the dimensionality was much higher and sustained for hundreds of milliseconds. These negative-control findings indicate that the results in Figure 2 specifically reflect the dimensionality of reliable stimulus-related signal.

Together, these findings show that natural images evoke rich multidimensional representations that rapidly evolve after stimulus onset and maintain their peak dimensionality for hundreds of milliseconds despite other ongoing sensory inputs. These findings further demonstrate that such multidimensional representations can be detected in large-scale EEG and MEG data and are specific to posterior channels over visual cortex.

### 2.2 Feature decoding tracks dimensionality

How do the dimensionality trends observed in Figure 2 relate to more conventional analyses of feature decoding? To address this question, we examined the decoding of both behavior-derived and DNN features, which have been a focus of previous work on object representations in EEG and MEG data [9, 41]. For the behavior-derived features, we examined a set of 66-dimensional embeddings that were learned in previous work to explain a large set of human similarity judgments on the objects from the THINGS database [24, 25]. For the DNN features, we examined a ResNet50 model trained on a large-scale dataset (YFCC15M) with a language alignment objective. Such language-aligned DNNs have been shown to be leading representational models of human visual cortex [12, 43]. We extracted activations from the first and last residual layers of this DNN to capture low- and high-level visual features.

For the decoding analyses, we fit linear regression models to predict behavioral and DNN features from EEG/MEG responses. For each time point, we fit these regression models using the training data and obtained decoding scores by computing the correlation between the actual and predicted feature values in held-out test data. Figure 2 shows the decoding score at each time point averaged across each set of behavioral and DNN features. Consistent with previous work [9, 41], each of these sets of features are significantly decodable within 100 ms after stimulus onset and remain significant for hundreds of additional milliseconds, while slowly decreasing in strength. As expected, we also observe different time courses for low- and high-level visual features. Specifically, we see that the decoding accuracy for early DNN features increases rapidly, reaching a peak around 100 ms, whereas the decoding accuracy for both behavioral and late DNN features rises more gradually, reaching its maximum between 200 and 400 ms. Analyses of other DNN layers are reported in Figure S2), and they show the expected temporal trends, with earlier layers tending to exhibit peak decoding accuracy at earlier time points.

When we compare the decoding results with the dimensionality trends in Figure 2a, we see that the strongest decoding accuracies are generally observed during the temporal window when neural dimensionality is sustained near its highest level. We also see a correspondence between the tails of the time courses for both decoding accuracy and dimensionality, which exhibit a similar pattern of slow decay. Together, these findings suggest that the dimensionality of stimulus-related neural activity may be a general indicator of expressivity, with times points of high dimensionality supporting the decoding of diverse behavioral and computational features. Furthermore, these findings suggest that neural dimensionality remains at its peak level even as the representations transform from low- to high-level features, as evidenced by the sustained dimensionality from approximately 100-400 ms despite ongoing changes in feature decoding.

### 2.3 Representations dynamically transform after stimulus onset

We next sought to understand how the underlying dimensions of these representations change over time. We began by selecting three example time points, and we visualized how the PCs from these time points evolve. As in the dimensionality analyses of Figure 2a, we learned the PC bases for each time point in the training data. However, instead of examining only the corresponding time point in the test data, we also examined cross-correlations with all other time points. As shown in Figure 3a-b, these analyses reveal that in both the EEG and MEG data, the PCs from early, middle, and late time points exhibit distinct trajectories, confirming that the representational content varies even during times when the overall dimensionality remains relatively stable. These analyses also show that the dynamics of representational changes are faster in the EEG data, which is consistent with the faster stimulus-presentation protocol in that study.

**Figure 3:**
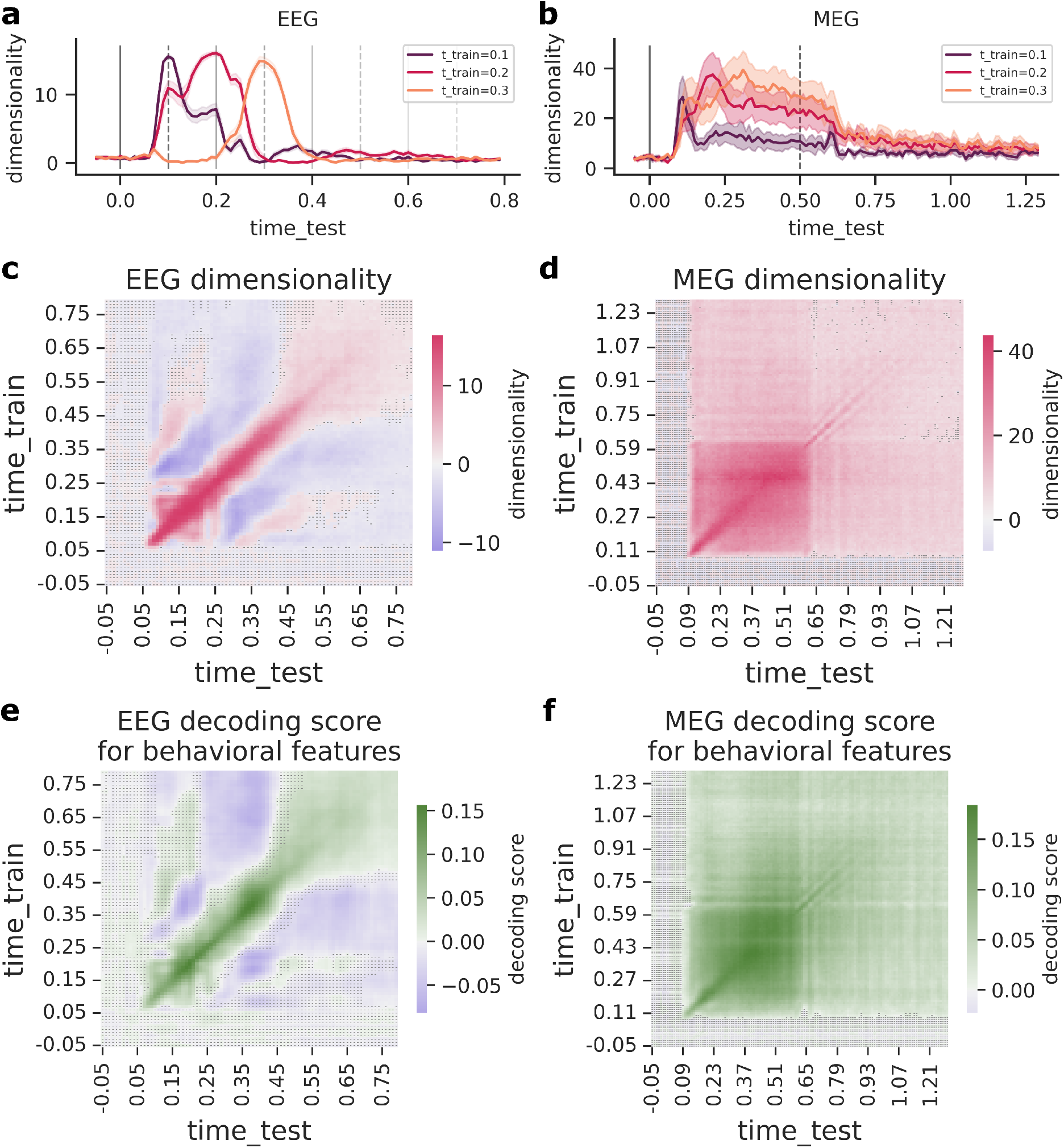
Temporal generalization of dimensionality and feature decoding. **(a), (b)** These plots illustrate how the PCs from three example time points evolve over time. At each time point relative to stimulus onset, these plots show the number of dimensions that have significant positive correlations with PCs learned from the selected example time points. The results show that in both the EEG and MEG data, representational PCs from earlier, intermediate, and later time points exhibit distinct temporal trajectories. They also show that the EEG data exhibit more rapid representational changes, consistent with the faster stimulus-presentation timing in the EEG experiment. **(c), (d)** These matrices show the number of dimensions shared across time points, with visualizations for both positively and negatively correlated dimensions. Positive dimensionality counts the number of dimensions along which two time points have significant positive correlations, while negative dimensionality counts the number of significant negative correlations. Each cell displays the greater value of the positive and negative dimensionality metrics. Shaded cells have no significant shared dimensions. Both datasets exhibit a band of strong positive dimensionality along the diagonal, and the EEG data additionally exhibit prominent oscillations from positive to negative dimensionality. **(e), (f)** These matrices show the average decoding scores for behavioral features, and they include all cross-time comparisons. Shaded cells have non-significant decoding scores. The trends in these matrices are broadly consistent with the dimensionality trends. Both datasets exhibit a band of strong positive decoding along the diagonal, and the EEG data additionally exhibit prominent oscillations from positive to negative decoding scores.

We next expanded these analyses to all time-by-time comparisons and generated matrices summarizing the number of correlated dimensions for each comparison (Fig.3c-d). When performing these analyses, we noticed that some cross-time comparisons in the EEG data exhibited predominantly negative correlations. To characterize this effect, we computed two dimensionality metrics: one for positive correlations and another for negative correlations. We used these two dimensionality metrics to generate the summary plots in Figure 3c-d, with each cell displaying the larger of the two metrics. Both the EEG and MEG data exhibit a band of high positive dimensionality along the diagonal, indicating that the multidimensional representations at each time point are reliable and at least partially distinct from other time points. However, the off-diagonal elements of these matrices show a clear difference across the EEG and MEG data. Specifically, in the EEG results, the dimensionality score fluctuates from large positive to large negative values, suggesting that all the dimensions of the representation are oscillating. We wondered if these oscillations might be related to the stimulus-presentation rate. To investigate this, we quantified how long it takes for the representations from a given time point to reach their peak negative dimensionality. We found that the modal time lag was approximately 200 milliseconds (Fig. S3), which corresponds to the time difference between consecutive stimuli, suggesting that the oscillations may be induced by the presentation rate. In contrast, negative correlations are absent from the MEG results. Instead, in the MEG data, the dimensionality scores for cross-time comparisons dissipate as the temporal difference increases, suggesting that the representations gradually decay when there are no intervening stimuli.

We also performed cross-time comparisons of the feature decoding models to see how they compare with the temporal patterns observed for dimensionality (Fig.3e-f). We focused on the average decoding accuracy of the behavior-derived features. As in the analyses in Figure 2c-d, we fit linear decoding models for each time point in the training data. However, instead of computing decoding scores using only the corresponding time points in the test data, we also computed decoding scores for every cross-time comparison, thus revealing the extent to which a decoder for a given time point generalizes to other times. The temporal patterns in these cross-time decoding results are similar to those in the dimensionality analyses in Figure 3c-d, with a band of strong positive decoding scores along the diagonal in both the EEG and MEG data. We also see oscillations in the off-diagonals in the EEG data, fluctuating from positive to negative decoding scores. Thus, these findings demonstrate that the dynamics of feature decoding track with the overall dynamics of the multidimensional neural representations.

Together, these cross-time analyses show that in both the EEG and MEG data, the content of the representations dynamically varies while the dimensionality is nonetheless sustained near its peak level for hundreds of milliseconds after stimulus onset. They further suggest that the structure of the representational dynamics is sensitive to the parameters of the stimulus-presentation timing, with faster and more consistent presentation rates potentially inducing strong oscillations in the stimulus-related responses.

### 2.4 Leading representational models do not fully account for neural dimensionality

We next sought to characterize the number of dimensions that are shared between the neural representations and the behavioral and DNN features. Our goal was to understand whether existing behavioral and DNN features fully account for the observed neural dimensions or whether the neural representations contain additional stimulus-related information that is not yet captured by leading theoretical models. To accomplish this, we performed a cross-decomposition analysis, which transforms both the neural responses and features to a shared orthogonal basis where their covariance is maximized [15, 22]. We used the same training and testing framework as the preceding analyses, with transformations learned on training data and applied to held-out test data. We computed a summary statistic of the shared dimensionality between the neural and feature representations by counting the number of shared dimensions whose reliability was above threshold in the test data. This is analogous to the neural dimensionality metric in Figure 2a-b.

The results are displayed in Figure 4. As expected, the temporal trends are broadly consistent with those observed for decoding accuracy (Fig. 2). We see an earlier peak for the dimensionality of low-level DNN features and later peaks for both high-level DNN features and behavioral features. Intriguingly, although the number of features in each set is generally larger than the number of neural channels (except when comparing behavioral embeddings and MEG data), we nonetheless find a substantial gap between the dimensionality of the neural representations themselves and their shared dimensionality with each feature set (Fig. 4). This holds true even when we combine the DNN and behavioral features in a single analysis, and it is consistent across all DNN layers examined (Fig. S4).

**Figure 4:**
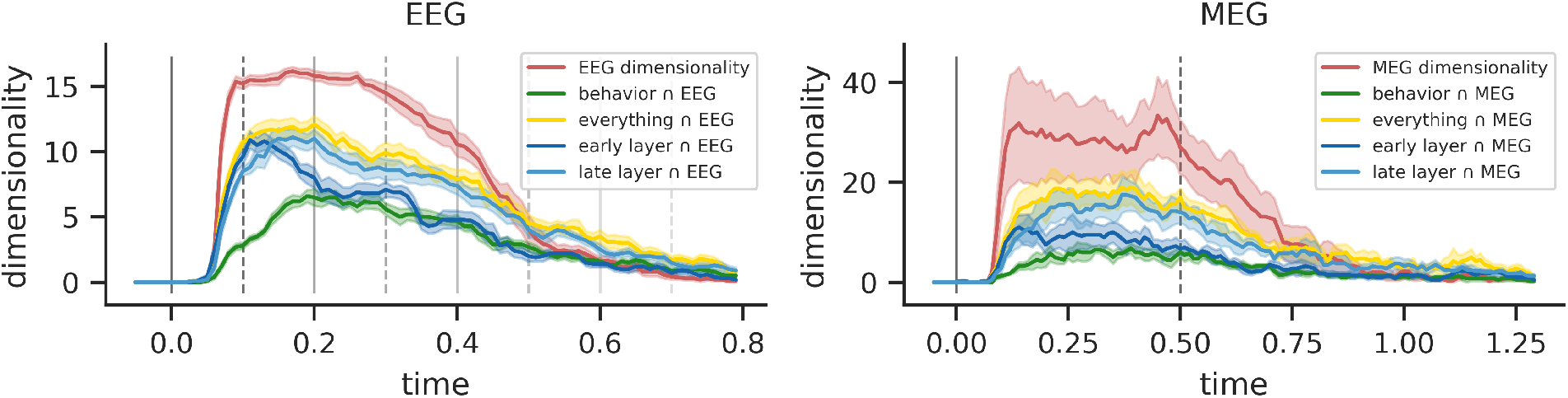
Temporal dynamics of shared dimensions between neural representations and visual features. These plots show the shared dimensionality between neural representations at each time point and the behavioral and DNN features. The dimensionality metric reflects the number of significant shared dimensions identified from cross-decomposition analyses comparing neural representations with behavioral and DNN features. The results labeled “everything” are for cross-decomposition analyses in which the behavioral and DNN features were combined. These plots also include the dimensionality of the neural representations themselves, which are also shown in 2. The vertical lines show the experimental setup of the two datasets, with solid lines indicating stimulus onsets and dashed lines indicating offsets. The error bars represent the standard error across subjects. These analyses reveal a substantial gap between the dimensionality of the neural representations themselves and the shared dimensionality with the behavioral and DNN features.

In sum, these findings show that leading computational and behavioral models do not yet account for the full complexity of stimulus representations that can be reliably detected in human EEG and MEG data.

### 2.5 Unexplained variance in neural representations contains behaviorally relevant information

We wondered whether the remaining unexplained variance in neural responses is perceptually relevant. Alternatively, it is possible that all the behaviorally relevant information might be contained within the subspace of the representations that is explained by the existing behavioral embeddings, which were derived from a large-scale dataset of human similarity judgments [24, 25]. As a case study, we focused on the EEG responses at 200 ms after stimulus onset, which is around the first peak of behavioral feature decoding. We examined the residual variance in the EEG responses to the test stimuli after regressing out the variance explained by the behavioral embeddings.

We began by visualizing these residual representations in a two-dimensional embedding using uniform manifold approximation and projection (UMAP) [35]. As expected, the residual representations do not exhibit readily interpretable clusters (Fig. 5a). This makes sense, since we have already regressed out the most salient factors driving behavioral similarity judgments. However, the remaining representations may nonetheless reflect perceptually relevant information, even if it is not easily interpretable.

**Figure 5:**
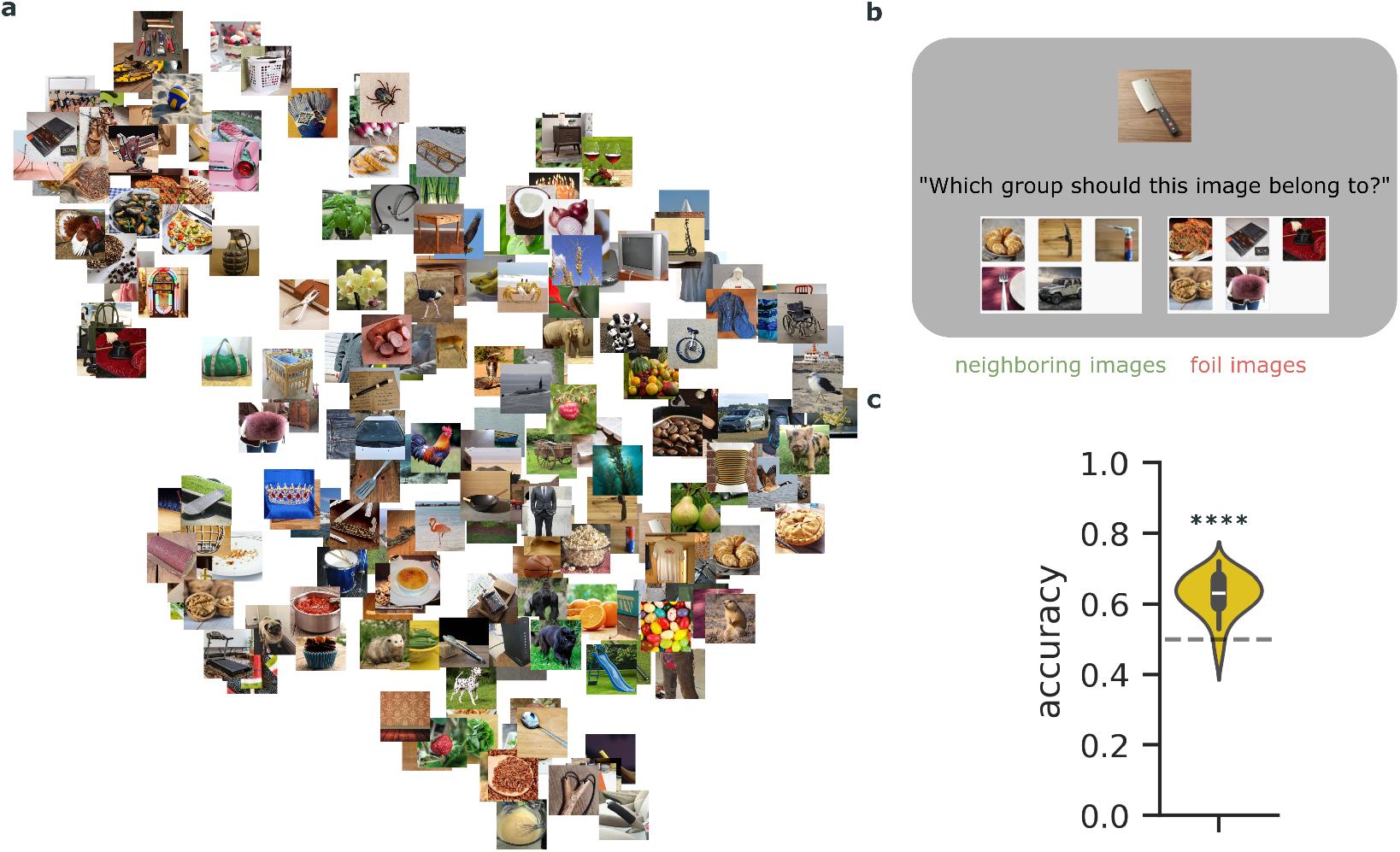
Behavioral relevance of residual variance in neural data after partitioning out embeddings learned from large-scale human similarity judgments. **(a)** This plot shows a UMAP embedding of the residual variance in EEG responses at 200 ms after partialling out the behavioral features learned from a large-scale dataset of human similarity judgments [24, 25]. As expected, the most salient interpretable structure in the data has been partialled out, but the remaining unexplained variance may nonetheless reflect perceptually relevant information. A behavioral experiment was designed to test this possibility. **(b)** In the behavioral experiment, participants (*N* = 50) chose which group the target images should belong to. The correct group includes five neighboring images with the closest Pearson distance to target image in the residual representations, while the foil cluster includes five random images that are distant from the target image. **(c)** This violin plot shows the distribution of accuracy scores across participants. Participant accuracy is significantly above chance, showing that the residual variance in the neural data reflects perceptually relevant information. **** *p* < 1*e* − 4

To test this possibility, we performed a behavioral experiment using the residual neural responses. We recruited online participants to perform an image-matching experiment, where the task was to chose which of two clusters a target image should belong to (Fig. 5c). The correct cluster includes five “neighboring” images based on Pearson distance (i.e., one minus correlation of the residual channel responses). The foil cluster includes five random images that are distant from the target. Participants’ accuracy was significantly above chance (*M* = 0.62, *SD* = 0.06, *t*(49) = 14.63, *p* < 1*e* − 4, Cohen’s d = 2.07; Fig. 5d). We also replicated these findings using the neural representations from another time point (100 ms) to confirm that they are not contingent on the specific time point selected (*M* = 0.57, *SD* = 0.05, *t*(49) = 9.34, *p* < 1*e* − 4, Cohen’s d = 1.32; Fig. S5).

Together these findings demonstrate that EEG responses to images encode rich behaviorally relevant information that extends beyond the most salient factors captured by conventional behavioral assessments of perceived similarity.

## 3 Discussion

We characterized the multidimensional dynamics of human EEG and MEG responses to natural object images using cross-validated matrix decomposition. Our findings reveal that the dimensionality of stimulus-related variance rapidly increases after stimulus onset and is sustained at its peak level for several hundred milliseconds. These neural dimensions are only partially explained by existing behavioral and computational models. Follow-up behavior experiments show that the remaining unexplained variance is behaviorally relevant, with participants reliably matching images based on their neural similarity. These results challenge prevailing assumptions about the inherent low-dimensional limitations of EEG and MEG recordings and reveal rich representational structure beyond what current models capture.

Our findings have implications for understanding the insights that can be gained from EEG and MEG data. Studies typically focus on leading components or predefined features in these recordings, potentially underestimating their representational richness [2, 4, 23, 8, 20]. Our results show that the stimulus-related information in EEG and MEG data extends well beyond the leading PCs. This suggests a remarkably rich encoding of visual information, consistent with recent work demonstrating complex multidimensional structure in visual cortex representations measured with fMRI [15, 22]. Moreover, temporally resolved recordings offer unique insights that fMRI studies cannot provide: we can track how the structure of the representations evolves millisecond by millisecond, revealing rapid multidimensional changes as visual processing unfolds. This temporal perspective complements the spatial richness of fMRI, demonstrating that both spatial and temporal resolution contribute distinct and valuable information about the dimensionality of neural codes.

Notably, the present analyses characterize dimensionality at individual time points. Future work could seek to reveal even richer multidimensional structure in EEG and MEG data by considering all time points together, as representations may utilize distinct dimensions at different moments throughout visual processing. However, simply concatenating responses across time and applying our decomposition analyses would conflate variance due to stimulus identity with variance due to temporal dynamics alone, making it difficult to identify which dimensions are actually stimulus-related. Alternately, one could concatenate all time points and channels and decompose these spatiotemporal response vectors, but the resulting latent dimensions would mix spatial and temporal information, making it difficult to examine temporal dynamics. Thus, while our work characterizes the multidimensional structure at each time point and the temporal evolution of this structure after stimulus onset, future studies could benefit from the development of new methods that capture the full range of orthogonal dimensions elicited in temporally resolved neural responses.

Our findings also reveal that there is a substantial gap between neural dimensionality and the information captured by representational models [24, 38, 29]. This gap persisted even when combining behavioral embeddings designed to capture human similarity judgments with different layer-wise embeddings of neural networks spanning early to late processing stages. This unexplained variance in EEG and MEG data suggests that current approaches do not yet account for the full complexity of neural representations. The behavioral relevance of this residual neural variance further demonstrates that unexplained dimensions contain meaningful perceptual structure. Participants could reliably match images based on residual neural patterns, showing that the human visual system encodes visual information along dimensions that are not yet fully captured by current models.

Our analyses used orthogonal decomposition methods to characterize the full spectrum of latent dimensions encoding stimulus related information in EEG and MEG data. Previous work has shown that representations from such time-resolved neural data can be mapped to those measured with spatially resolved data, like fMRI, allowing researchers to characterize how stimuli are represented over both time and space [8]. However, these previous methods relied on the use of RSA, which is largely driven by the leading PCs of neural responses and may overlook other meaningful but low-variance dimensions [15, 22]. In contrast, our cross-decomposition method is sensitive to the full spectrum of latent dimensions in neural representations. In future work, this cross-decomposition method could be adapted to identify shared dimensions across recording modalities, thus potentially revealing a more detailed picture of the spatiotemporal structure underlying dynamic human brain responses.

Our findings show that, depending on experimental conditions, image representations can exhibit more complex temporal structure than current theories emphasize. Specifically, previous work has largely focused on the distinct temporal profiles observed for different visual features [30, 4, 8, 9, 10, 20, 41]. We replicate some of these previous findings by showing earlier decoding of low-level features and later decoding of high-level features. However, our findings in the EEG data additionally reveal an unexpected oscillatory pattern in the stimulus encoding, whereby the entire spectrum of representational dimensions oscillates. This means that the stimulus information encoded at one time point is also observed at later time points but with the opposite polarity. We suspect that this transient oscillatory pattern could be induced by the rapid and regular temporal structure of stimulus presentation in the EEG experiment. However, even if this explanation is correct, it raises important questions for future work. First, it remains unknown what functional relevance these oscillations might have, if any. One possibility is that they are related to oscillatory mechanisms of visual attention [13]. Second, it highlights the possibility that conventional representational analyses, such as feature decoding, could easily overlook major representational transformations if they are applied to each time point independently or on temporally averaged data.

Our work reveals that time-resolved neural responses to natural images contain rich multidimensional structure that approaches the capacity of the recording modalities. This structure cannot be fully explained by current representational models, whether derived from similarity judgments or deep neural networks, yet it contains behaviorally relevant information. By revealing rich structure in modalities often considered noisy or limited, our results open new avenues for investigating the neural basis of vision. Together, these findings highlight the importance of going beyond the leading dimensions of neural activity and understanding dynamic human brain responses in their full complexity.

## 4 Materials and Methods

### 4.1 Datasets

#### 4.1.1 Dynamic neural data

We examined neural responses to natural object images from the THINGS database, which contains images from 1,854 diverse object categories [27]. We analyzed two large-scale publicly available datasets of dynamic neural responses to these images. The EEG dataset (“THINGS-EEG2”) includes ten subjects who viewed THINGS images and determined if a target image showed up in a given sequence [16]. Each stimulus was presented for 100 ms, followed by 100 ms of fixation, after which the next stimulus was shown. The training set includes four repetitions of responses to ten unique images per 1,654 concepts; the test set includes 80 repetitions of responses to one image per 200 concepts, which do not overlap with the concepts in the training set. The epoched data range from 200 ms before stimulus onset to 800 ms after stimulus onset. We used the data from 17 channels in the occipital and parietal lobes, selected and preprocessed by [16].

The MEG dataset (“THINGS-MEG”) [25] includes four subjects who viewed THINGS images and responded whenever they detected an oddball synthetic image. Each stimulus was presented for 500 ms, followed by 1 second fixation, with ±200 ms of jitter, after which the next stimulus was shown. The training set includes 12 unique images per 1,854 concepts, each presented once; the test set includes one image per 200 concepts, each presented 12 times. We sought to match the preprocessing procedures across the EEG and MEG datasets as much as possible, so we performed our own preprocessing on the raw MEG data with a few differences relative the preprocessing performed by the original authors. These differences include bandpass filtering the raw data for each run between 0.1 and 100 Hz, baseline correcting the epoched data by subtracting the mean of the data during baseline 100 ms before stimulus onset, and downsampling to 100 Hz. Because differences in stimulus timing across the EEG and MEG experiments, we used the same epoch time window for the MEG data as the original authors [25], which ranged from –100 ms to 1300 ms relative to stimulus onset. Our main analyses used the 80 channels from the occipital and parietal lobes.

#### 4.1.2 Behavioral embeddings

We analyzed 66-dimensional sparse positive object similarity embeddings (SPoSE) for the 1,854 object categories. These embeddings were learned in previous work to explain 4.7 million trials of human odd-one-out judgments on the THINGS images [24, 25].

#### 4.1.3 Neural network feature maps

We extracted feature maps from OpenCLIP ResNet50, a state-of-the-art vision neural network trained on the YFCC15M dataset [38, 29]. We extracted activations from the first and last residual units, using the Identity layer following the middle ReLU. We reduced the dimensionality of these DNN activations using sparse random projections, with the the number of projections determined via the Johnson-Lindenstrauss lemma, resulting in 8,336 dimensions.

### 4.2 Linear mapping methods

#### 4.2.1 Feature decoding

We used ridge regression to predict features of interest from neural responses at each time point. For all analyses, we used a ridge penalty of *α* = .01. We confirmed that our findings are not strongly contingent on the specific value of the ridge penalty. For each subject and time point, regression weights were learned using the training set neural responses averaged across repetitions. Decoding scores were computed by taking the Pearson correlation between the predicted and actual feature values in the test set, with statistical significance assessed using cluster-based permutation tests [34]. When computing decoding scores for the temporal generalization analyses, the regression weights learned at a given time point were applied to the test-set neural responses at all other time points.

#### 4.2.2 Neural dimensionality

We assessed the dimensionality of stimulus-related variance in neural responses at each time point [40, 15, 22]. For the training-set, we computed the average responses across all repetitions, *X*_train_; for the test set, we randomly split the repetitions into two halves and computed the average responses within each half, 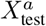 and 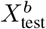. We first fit PC eigenvectors on the training set:

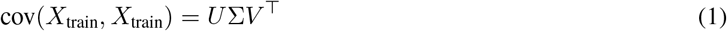

We then projected the test-set responses onto the learned eigenvectors, 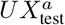and 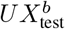. At each rank, we calculated reliability by taking the Pearson correlation across the test-set splits, with statistical significance assessed using cluster-based permutation tests [34]. We computed a summary statistic of the representational dimensionality at each time point by counting the number of significant dimensions. For temporal generalization analyses, the eigenvectors learned at a given time point were applied to test-set neural responses at the same time point and at all other time points. Correlations were then computed across time points rather than across stimulus repetitions.

#### 4.2.3 Shared-space dimensionality

We computed the shared dimensionality between neural responses and representational features using a similar approach to the neural dimensionality analyses. We computed the average responses across all repetitions, *X*_train_ and *X*_test_. Given the neural responses and a set of features, *Y*_train_ and *Y*_test_, we performed singular vector decomposition (SVD):

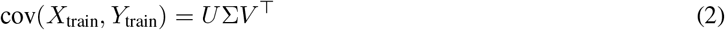

We then projected the test-set representations onto the shared space, *UX*_test_ and *Y*_test_*V*. Again, we calculated the Pearson correlation between the projections at each rank and defined the shared dimensionality as the number of dimensions significantly above the null distribution using cluster-based permutation tests [34].

### 4.3 Behavioral experiment

We conducted an image matching experiment to assess if reliable neural signals unexplained by existing behavioral embeddings are also behaviorally meaningful. We chose to focus on the EEG data at time point 200 ms, approximately where the shared dimensionality between EEG data and behavioral embeddings peaks. We first trained a ridge regression (*α* = .01) to predict the train-set neural data, averaged across all repetitions, from the behavioral embeddings. We then computed the residual variance in the test set by subtracting the predicted values from the real test-set responses, averaged across all repetitions. We calculated the pair-wise Pearson distance (one minus Pearson correlation coefficient) between the residual responses to the test stimuli for each subject and averaged these values across subjects.

In the experiment, participants (*N* = 50) were presented with a target image on top and two groups of five images below. They were asked which group the target image should belong to. The correct group contains five images with the closest Pearson distance to the target image, and the foil group contains five images that were randomly selected from images whose distance to the target was above a fixed threshold (we used the 0.8 quantile of all pairwise distances in the first experiment, and we relaxed this to the 0.65 quantile in the replication experiment on EEG responses at 100 ms). Experimental trials were created for all 200 test images. The correct image group appeared randomly on the left or right side of the display across trials. The participants first completed a practice trial with the correct group being all the same image as the target images. Then they proceeded to complete a sequence of 100 randomly sampled experiment trials and ten catch trials in the same format as the practice trial. Experimental trials were sampled such that across sets of two participants, all 200 trials would be shown (i.e., for every 100 trials selected for one participant, the remaining 100 would be shown to another participant).

## 5 Data Availability

The EEG dataset (THINGS-EEG2) is publicly available at [16, 17]. The MEG dataset (THINGS-MEG) is publicly available at [25, 26]. Anonymized behavioral data collected in this study are available at [6].

## 6 Code Availability

The code for reproducing all analyses and figures in this manuscript is available at [7].

## 8 Supplementary Materials

**Figure S1:**
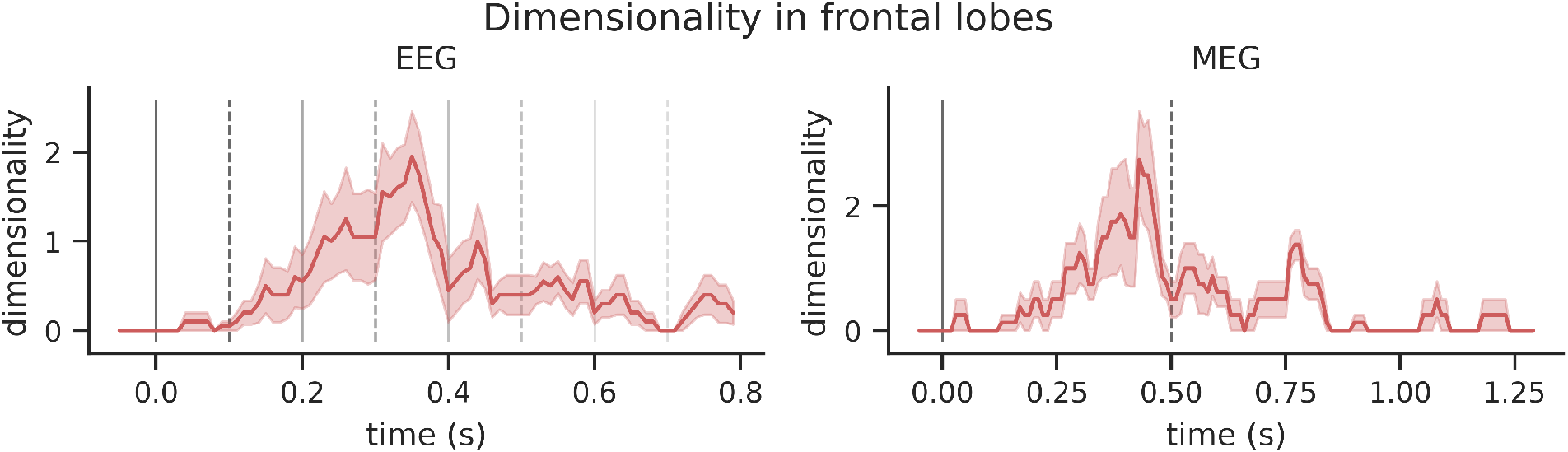
Temporal dynamics of neural dimensionality in frontal channels. These plots show the dimensionality of stimulus-related variance in channels over the frontal lobes in the EEG and MEG datasets. The error bars represent the standard error across subjects. The vertical lines show the stimulus-presentation timing of the two datasets, with solid lines indicating stimulus onsets and dashed lines indicating offsets. The dimensionality of these frontal channels is much lower and more transient than the dimensionality observed over posterior visual regions in Figure 2.

**Figure S2:**
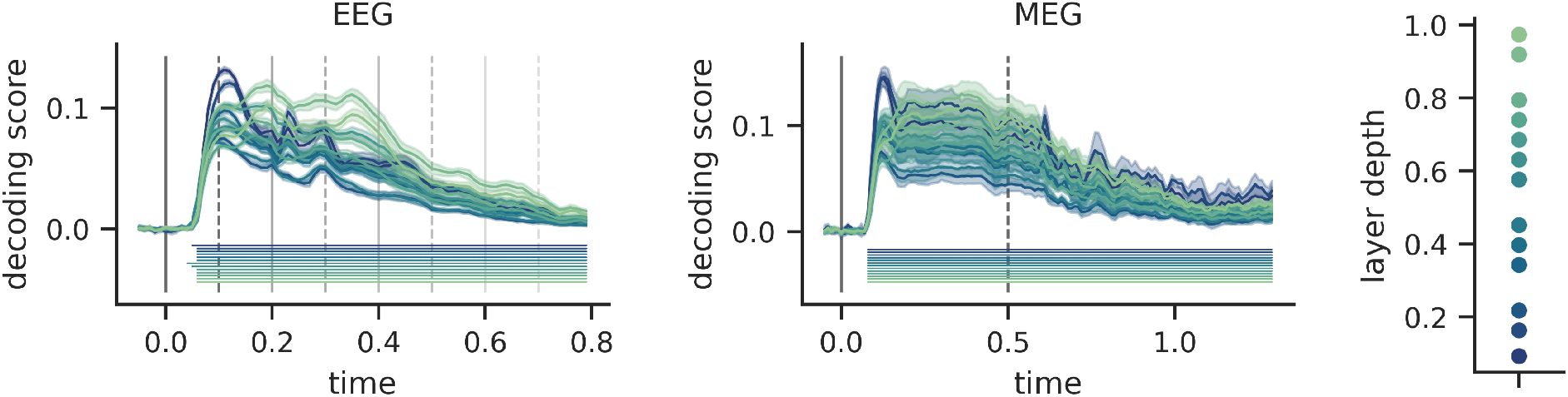
Temporal dynamics of decoding scores for DNN features across layer depths. These plots show the average decoding scores for DNN features at different layers. The vertical lines show the stimulus-presentation timing of the two datasets, with solid lines indicating stimulus onsets and dashed lines indicating offsets. The error bars represent the standard error of feature-averaged means across subjects. The right subplot shows the relative layer depths. Features from all layers can be decoded from the neural responses. Further, these decoding time courses exhibit a trend consistent with previous findings in which decoding peaks earlier for lower-level features and later for higher-level features [30, 4, 8, 9, 10, 20]. Horizontal lines under the decoding time courses indicate time windows of significant decoding (*p* < 0.05, cluster-based permutation test).

**Figure S3:**
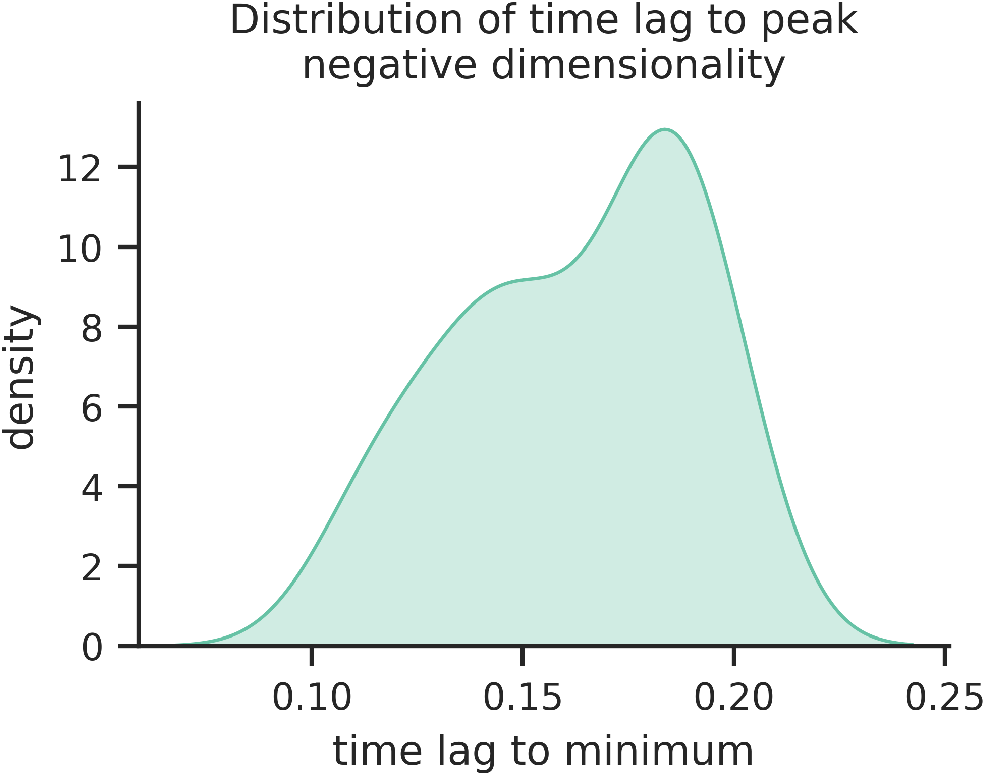
Distribution of time lag to peak negative dimensionality in EEG temporal generalization. For each training time point between 100 and 400 ms post stimulus onset in the EEG data, we identified the test time at which negative dimensionality reached its highest magnitude, and we computed the absolute time lag to this peak. The range of training time points examined here corresponds to the window of high positive dimensionality in Figure 2a. The distribution of these time lags shows a mode at approximately 200 ms, corresponding to the inter-stimulus interval in the EEG experiment (100 ms stimulus presentation followed by 100 ms fixation). This result suggests that the oscillatory sign flips observed in the off-diagonal bands of the EEG temporal generalization matrix (Fig. 3c) may be driven by the stimulus presentation rate.

**Figure S4:**
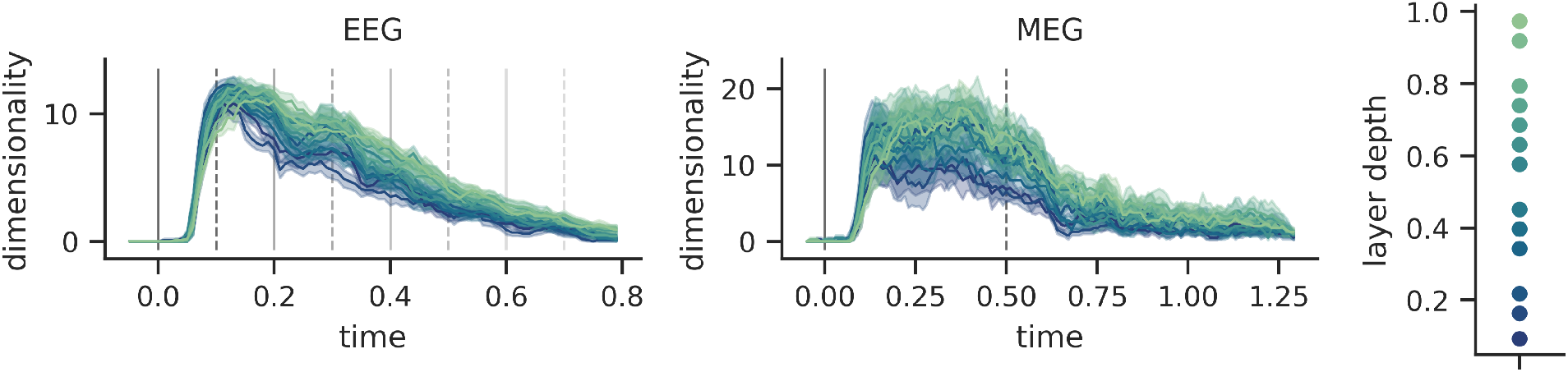
Temporal dynamics of shared dimensions between neural representations and DNN features across layer depths. These plots show the shared dimensionality between neural representations and DNN features at different layers. The dimensionality metric reflects the number of significant shared dimensions identified from cross-decomposition analyses comparing neural representations with DNN features. The vertical lines show the experimental setup of the two datasets, with solid lines indicating stimulus onsets and dashed lines indicating offsets. The error bars represent the standard error across subjects. The right subplot shows the relative layer depths. Consistent with the layer-wise decoding scores in Figure S2, these plots exhibit a trend in which shared dimensionality peaks earlier for lower-level features and later for higher-level features.

**Figure S5:**
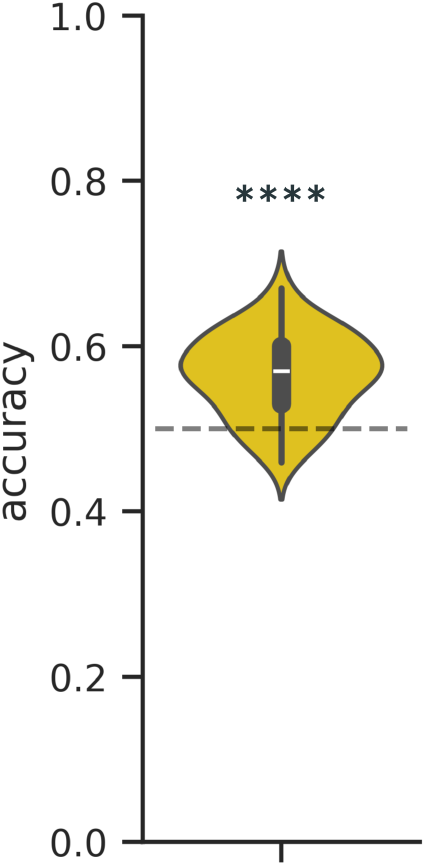
Behavioral performance on the image-matching experiment based on residual representation from the EEG data at time point 100 ms. The behavioral finding in Figure 5 was replicated using EEG responses from a different time point (100 ms) (*N* = 50 participants). This violin plot shows the distribution of accuracy scores across participants. Participant accuracy is significantly above chance, showing that the residual variance in the neural data reflects perceptually relevant information. **** *p* < 1*e* − 4

